# When communication fails, physical effort increases but not to greater effect

**DOI:** 10.64898/2026.07.06.736716

**Authors:** Šárka Kadavá, Wim Pouw, Susanne Fuchs, Judith Holler, Aleksandra Ćwiek

## Abstract

Successful communication depends not just on what we say but also on how we repair it when understanding breaks down. People routinely respond to misunderstanding by gesturing more or speaking louder. Yet whether these responses actually reflect measurable bodily effort, and whether it helps communication, has not yet been directly tested. Extending the hypo- and hyper-articulation accounts to whole-body multimodal communication we ask: does regaining understanding require increased physical effort, and does that effort improve communicative outcomes? In a pre-registered biomechanical study, 61 pairs of participants conveyed concepts without conventionalized language, using gestures, vocalizations, or both, and repaired failed attempts. We measured communicative effort by capturing arm torque change (upper-limb effort), vocal amplitude (vocal effort), and postural adjustments (postural effort). Using Bayesian hierarchical models, we show that repair increases physical effort across all three measurements, but that performers communicate more over time rather than more forcefully at any particular moment in time. Upper-limb effort scale with the degree of misunderstanding but effort overall does not predict whether communication succeeds. We suggest that communicative effort on the level we measured here functions as an index of continued commitment rather than a mechanism for repairing misunderstanding.

## Main

Communication is like a dance; it takes two partners, requires continuous effort, and when one stumbles, the other may work harder to keep going. When communication fails, partners may not only repeat themselves, but also speak louder or gesture more extensively, investing more physical effort to sustain the exchange. Yet although this bodily investment is central to everyday multimodal communication, its physical dimension remains largely unexplored. The current pre-registered study investigates whether communicative modulations people make in response to misunderstanding are associated with a measurable increase in physical effort and what that cost reveals about the body’s role in keeping communication alive.

Across locomotion, arm-reaching, and decision-making, actions emerge from the continuous negotiation of metabolic, computational, cognitive, and biomechanical constraints (Bernstein, 1967; Ferrer-i-Cancho et al., 2013; Kelso, 1995; Kool et al., 2010; McAllister et al., 2025; Rosen, 1967; Tversky & Kahneman, 1974; Wong et al., 2021). This is similarly true for language: the way that people speak and structure their utterances reflects effort, particularly the optimization toward low-cost solutions (Gibson et al., 2019; Levshina, 2022; Levshina & Moran, 2021; Zipf, 1949). In linguistics, effort has been operationalized primarily in terms of message length and information structure. Short, frequent forms require less articulation and place fewer demands on processing (Zipf, 1949), and the principle of effort optimization has been argued to shape lexical frequency, word order, syntactic structure, sound duration, and articulatory detail across languages (Aylett & Turk, 2004; Bell et al., 2009; Futrell et al., 2015; Haspelmath, 2021). These regularities, often explained in information-theoretic terms, may themselves have a physical basis (Torre et al., 2019). In speech sciences, physical effort has been used to explain why vocal targets are often not fully realized, resulting in phonetic reductions (Kirchner, 2001; Lindblom, 1990). What these traditions have in common is that they have focused almost exclusively on the structure and production of spoken language, with a few exceptions extending these accounts to sign (Napoli et al., 2014; Slonimska et al., 2022).

Yet, language is fundamentally multimodal, distributed across, e.g. hand gestures, eye gaze, facial expressions, head nods, postural sway (Emmendorfer & Holler, 2025; Nota et al., 2021; Shockley et al., 2003; ter Bekke et al., 2024), and each of these bodily signals emerges from movement shaped by the constraints of human physiology (Kadavá et al., accepted). Speech is shaped by breathing (Fuchs & Rochet-Capellan, 2021) and the biomechanics of the tongue and mandible (MacNeilage & Davis, 2000); head and arm gestures involve substantial mass that interacts with body systems crucial for vocalization, such as the larynx (Ćwiek & Fuchs, 2025) and the lungs (Pouw et al., 2020); and the forces these movements generate are integral to their communicative function (Krebs et al., 2025). Finally, posture provides the stable foundation on which all these bodily signals depend (Massion, 1992). Movement is therefore a primary component of communication, and like all movement, it carries a physical cost.

What distinguishes communicative movement from other types of movement (e.g. reaching) is that it is a signal produced for someone: an act of skilful intersubjective coordination (Di Paolo et al., 2018; Fusaroli et al., 2014). Lindblom’s (Lindblom, 1990) H&H Theory captured this as a trade-off between biological (optimizing) processes and environmental constraints, positioning behavior on a continuum between hypo- and hyper-articulation. Most prior work on hyper-articulation has used visual or kinematic correlates across communicative contexts such as infant-directed speech and gesture (Biersack et al., 2005; Brand & Shallcross, 2008; Rohlfing et al., 2022; Song et al., 2010); pantomime (Żywiczyński et al., 2023); speech addressed to non-native or hearing-impaired interlocutors (Beechey et al., 2020; Knoll et al., 2015; Smith, 2007); and communication in background noise (Dohen & Roustan, 2017; Garnier & Henrich, 2014; Trujillo et al., 2021). A particularly revealing window into how the hypo- vs. hyper-articulation trade-off is realized in social interaction comes from repair, i.e. the moments when understanding breaks down and must be restored (Dingemanse et al., 16. 9. 2015; Schegloff et al., 1977). Unlike noise or perceptual impairment, misunderstanding imposes an epistemic constraint: the signal was received, but its meaning is not shared. Speakers add more detail to their gestures when they recognize a lack of shared understanding (Hilliard & Cook, 2016), produce more submovements, and use more orthographic characters when attempting to regain it (Rasenberg et al., 2022). Addressee’s feedback can also cause gestures to increase in rate (Hoetjes et al., 2015) and become more precisely articulated than their previous occurrence (Holler & Wilkin, 2011). The opposite applies when common ground is established: gestures become shorter (Holler et al., 2022), smaller in amplitude and less precise (Galati & Brennan, 2014; Gerwing & Bavelas, 2004), or less frequent (Holler & Stevens, 2007; Parrill, 2010; but see Holler & Bavelas, 2017 for exception). However, this body of work has typically framed these adaptations in terms of clarity, informativity, or perceptual robustness rather than physical effort, and has relied on qualitative observations or surface kinematics and acoustic proxies rather than direct biomechanical measurements. Therefore we are not able to determine whether reported changes are only reorganizations of the communicative system, or whether they (also) recruit more bodily resources. This is a fundamental gap, because evolutionary accounts of communication rest on the assumption that signals are costly: that animals communicate with minimal effort when conditions allow, and with greater effort when they must (Hex et al., 2026; Zahavi et al., 1999). Without grounding this cost in resources that are actually recruited by the body the trade-off at the heart of H&H Theory remains an assumption rather than a demonstrated mechanism.

In the current study, we test whether the modulations that people make in their communicative behavior to regain understanding are associated with an increase in physical effort. We define physical effort as an energetic cost of an action (Halperin & Vigotsky, 2024) associated with the enhancement of (a feature of) an articulator (Lindblom, 1990), and treat increased effortfulness as a shift from a low-cost baseline toward hyper-articulation. To measure this, we rely on biomechanical signals which are estimates of the mechanical work performed by the body (i.e. the forces used to displace body segments) and which therefore offer a more direct proxy of metabolic cost than kinematic or visual correlates predominantly used in prior work. Previous studies have relied on measures such as gesture size, gesture frequency or number of submovements – which capture the spatial and temporal profile of movement but are insensitive to the forces required to produce it. Because force depends on the mass of the moved segment and its acceleration rather than on spatial amplitude alone, two movements that look similar kinematically can differ substantially in the underlying physical effort, and vice versa (Hagen & Valero-Cuevas, 2017). Specifically, we combine the acoustic amplitude envelope, which reflects subglottal pressure and respiratory engagement (Fuchs & Rochet-Capellan, 2021; Pouw et al., 2023; Zhang, 2016), with two biomechanical measurements: joint torque change for upper-limb movement (Uno et al., 1989) and change in center of pressure (COPc) for postural adjustments (Winter, 1995). Both have been shown to predict metabolic cost (Mohammadzadeh Gonabadi et al., 2024) and, unlike kinematic proxies, account for the masses being moved and the dynamic demands of maintaining balance.

We further distinguish two dimensions of physical effort. An increase in effort could be realized either by intensifying a given action – striking harder, speaking louder, exerting greater peak force (henceforth ‘instantaneous effort’) – or by extending the action over a longer window without necessarily increasing its peak intensity (henceforth ‘cumulative effort’). If effort is concentrated in a single moment, peak intensity and total cost are essentially the same thing. But once effort is spread out across several moments or a longer stretch of time (e.g. several gesture strokes), the two can come apart: one could, for instance, exert the same modest force at each moment but sustain it for longer, raising the total cost without raising the peak at any single point. Distinguishing instantaneous from cumulative effort therefore allows us to ask not only whether physical effort increases under misunderstanding, but whether that increase is concentrated locally or distributed across time.

We test two pre-registered hypotheses (H1, H2). In our referential task design (Fig. 1a), participants convey concepts without conventionalized language, using gestures, vocalizations, or both. Each failed guess from their partner triggers an obligatory correction attempt. Based on prior research (Lindblom, 1990), we expect these correction attempts to result in hyper-articulated behavior. Thus, we predict that *correction recruits more physical effort than baseline performance* (H1). Further, because interlocutors are known to adapt their multimodal signals depending on shared knowledge with their partners (Gerwing & Bavelas, 2004; Hilliard & Cook, 2016; Holler & Wilkin, 2011; Levshina, 2022), we assume that the guesser’s answer provides the performer with possibility to assess the degree of common ground and this with the opportunity to act on it, taking into consideration how far this answer is from the target meaning (for instance, by enhancing communicative movements). Concretely, we predict that *a higher degree of misunderstanding recruits more physical effort from the performer* (H2). Because the communicative modality available for use in the different experimental conditions determines which articulators are available for elaboration, we additionally examine how effort levels and communicative outcomes differ across unimodal and multimodal conditions, asking whether having access to both gesture and voice saves effort. In an exploratory analysis that was not pre-registered, we further ask whether the effort invested in repair translates into better communicative outcomes, a question that, to our knowledge, has not yet been systematically examined. Across the three modalities, we find that in communicative repair, performers recruit cumulative but not greater instantaneous effort, a pattern that reflects elaboration of the message rather than amplification of its parts. This distinction reframes effortful repair as the production of more communicative material rather than delivering it more intensely, with implications for how the body coordinates to sustain mutual understanding.

**Fig. 1:**
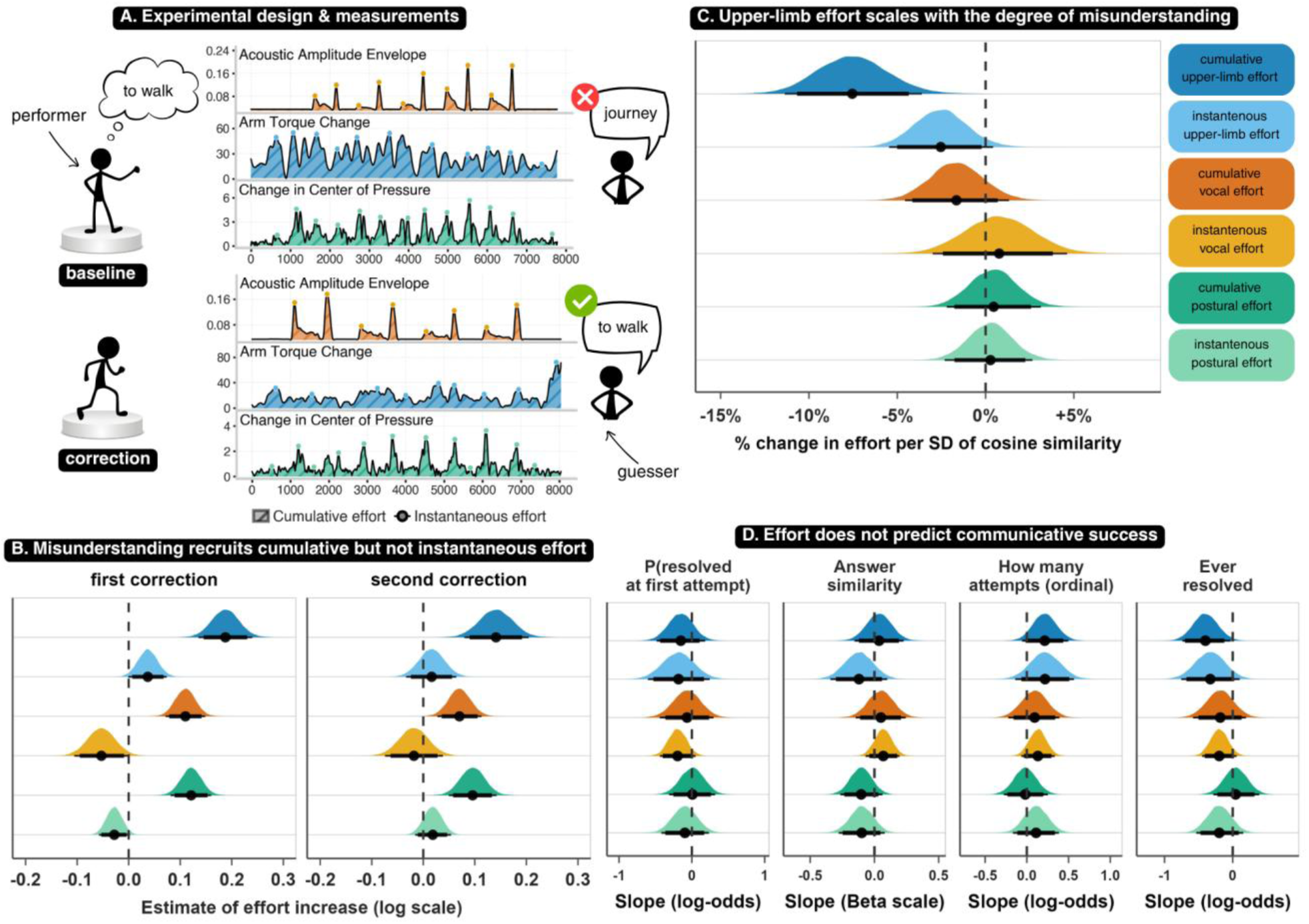
Task design and empirical results. (A) Experimental design and measurements. A performer expresses a target concept to a guesser under one of three modality conditions (gesture-only, vocal-only, or multimodal); after an incorrect response, the performer produces a correction, up to two corrections per concept (baseline guess ‘journey’, incorrect; correction guess ‘to walk’, correct). We extract three effort signals: torque change of the arms (upper-limb effort), acoustic amplitude envelope (vocal effort), and change in center of pressure (postural effort), each summarized as cumulative effort (area under the curve) and instantaneous effort (mean of detected peaks). These give the six effort variables used throughout. (B) Posterior distribution of effort increase across corrections, on the log scale: first correction (left panel) is relative to baseline, and second correction (right panel) is relative to first correction. Dashed line at 0: no change; point: posterior median; thick and thin lines: 89% and 95% highest density intervals (HDI). (C) Posterior distribution of effort increase per 1 SD of cosine similarity between the answer and the target meaning (%, see Methods); higher similarity reflects less misunderstanding. Conventions as in B. (D) Posterior slopes relating the six effort variables to four measures of communicative success, each a separate model (see Methods): probability of understanding at the first-attempt (log-odds); answer similarity to target (Beta scale); understanding across attempts (ordinal: first attempt < first correction < second correction < unresolved; log-odds); and whether ever resolved (log-odds). Conventions as in B.

## Results

### Misunderstanding recruits cumulative but not instantaneous effort

As predicted, participants invested more effort when their partners misunderstood them but the pattern was more specific than a simple mobilization of all available resources. Here we report the main effects of interest, a full report of the results is available in Supplementary Material (section F). Across all investigated dimensions of effort – upper-limb, vocal, and postural – behavior following an incorrect guess was more effortful in cumulative but not instantaneous terms (Fig. 1b). Upper-limb effort showed the largest enhancement: cumulative torque change increased by 20.6% at the first correction compared to baseline (95% CrI: [14.5%, 27.1%]) and by 15.2% at the second correction compared to first correction (95% CrI: [8.1%, 22.6%]). Vocal and postural effort followed the same pattern at roughly half the magnitude. Cumulative vocal effort increased by 11.6% at the first correction (95% CrI: [7.4%, 16.0%]) and by 7.3% at the second (95% CrI: [2.8%, 12.0%]), and cumulative postural effort increased by 12.9% at the first correction (95% CrI: [8.5%, 17.4%]) and by 10.0% at the second (95% CrI: [5.0%, 15.2%]).

To test whether these cumulative increases reflected more movement or larger movements, we refitted the models with trial duration as a covariate. Adding duration substantially attenuated the correction effect (cf. Supplementary Material section F, Table 9). The first-correction effect on the upper-limb effort remained credibly positive, but decreased from 20.6% to 7.0% (95% CrI: [2.6%, 11.7%]), suggesting that almost two-thirds of the increase in upper-limb effort can be attributed to accumulation of more, rather than larger, arm movement. The second-correction effect on upper-limb effort and both correction effects for vocal and postural effort, were fully accounted for by duration: with duration in the model, none remained credibly different from zero.

None of the instantaneous measures showed a similar credible pattern of increase: at the populational level, the instantaneous vocal, upper-limb, and postural effort remained unchanged following a misunderstanding. In an exploratory analysis, however, a model including the interaction between correction and each performer’s average baseline (i.e. first performance) effort showed that the first-correction effect is credibly influenced by the amount of effort in the baseline performance, and can even lead in the opposite direction (see Supplementary Material section F, Fig. 4). Specifically, for instantaneous arm torque change, the interaction with baseline effort was strongly negative: performers with low average baseline effort (−2 SD from the mean) increased their first correction by 17.2% (95% CrI: [8.5%, 26.6%]), while those with high baseline effort (+2 SD) decreased it by 9.4% (95% CrI: [-16.4%, -1.8%]). Instantaneous vocal effort showed the same dependence: low-baseline performers (-2 SD) increased their first correction by 18.6% (95% CrI: [6.7%, 32.0%]), while high-baseline performers (+2 SD) decreased it by 24.6% (95% CrI: [-32.5%, - 15.7%]). For postural effort, the pattern was weaker: high-baseline performers decreased their first correction by 11.0% (95% CrI: [-16.6%, -5.1%]), while low-baseline performers showed no credible change (β = 6.5%, 95% CrI: [-0.2%, 13.7%]). Instantaneous effort at correction was therefore not uniformly recruited; whether performers scaled up the instantaneous effort depended on how effortful their baseline performance already was. This pattern warrants caution for two reasons. First, the baseline-by-correction interaction is susceptible to regression to the mean: participants with ‘extreme’ behavior will tend to regress toward the group mean at subsequent timepoints, which could artifactually produce the observed sign reversal. Second, high-baseline performers may face ceiling constraints on instantaneous effort, making decrease more likely by default. Because the present analyses cannot distinguish between statistical artifacts and behavioral patterns, we do not interpret it substantively.

We also asked how baseline effort levels differed across modality conditions, since the available articulators differ between them. The modality shaped effort, but in measure-specific ways. Vocal effort was substantially higher in the vocal-only than in the multimodal condition, both cumulatively (36.0% higher, 95% CrI [19.6%, 54.6%]) and instantaneously (58.5% higher, 95% CrI [34.0%, 87.4%]), implying that vocal effort takes on more of the communicative burden when the hands are unavailable. The effect of modality on postural effort is reversed in direction: conditions involving bodily movement elicited far greater postural adjustments than the vocal-only condition, both cumulatively (gesture-only: 313.3% higher, 95% CrI: [212.8%, 444.2%]; multimodal: 344.3% higher, 95% CrI: [235.4%, 486.9%]) and instantaneously (gesture-only: 147.9% higher, 95% CrI: [103.3%, 203.3%], multimodal: 188.7% higher, 95% CrI: [135.7%, 254.4%]). Gesture-only and multimodal conditions did not differ in cumulative postural effort, although multimodal exceeded gesture-only for instantaneous postural effort, suggesting that coordinating two modalities involves bursts of postural activity without increasing total postural load. For upper-limb effort, we found no reliable average difference between conditions once between-participant and between-concept variability was modelled. The gesture-only vs. multimodal contrast was positive for cumulative effort but its interval included zero (5.6%, 95% CrI [-1.7%, 13.5%]). Substantial random slope variance indicated that who is communicating and which concept is conveyed matter more for upper-limb effort than the modality condition itself.

To summarize, while cumulative effort increased reliably across corrections for all three measures, instantaneous effort did not. Correction therefore imposes a clear cumulative cost on performers without recruiting greater peak intensity per movement or per vocalization. This pattern is consistent with a redistribution of effort across longer time intervals and more movement units rather than a concentrated scaling-up of force or amplitude, an interpretation we develop in the Discussion. The consistency of these increases across three independent measures of bodily effort supports that correction imposes a substantive physical cost on the performer, supporting H1 for cumulative, though not instantaneous, effort.

### Upper-limb effort scales with the degree of misunderstanding

We found partial support for H2 (Fig. 2c). Cumulative upper-limb effort scaled credibly with the degree of misunderstanding, operationalized as the cosine similarity between the guesser’s previous answer and the target meaning (lower similarity indicates greater misunderstanding). Each 1-SD increase in cosine similarity was associated with 7.5% reduction in cumulative arm torque change (95% CrI: [-11.3%, -3.5%]); equivalently, performers invested more cumulative upper-limb effort when the previous answer was further from the target. Instantaneous arm torque change ran in the same direction, but its estimated interval included zero (β = -2.5%, 95% CrI: [-5.4%, 0.5%]). In contrast, neither cumulative nor instantaneous vocal effort showed a credible relationship with cosine similarity, and the same was true for both postural measures.

Cumulative upper-limb effort therefore adapts reliably to the amount of repair work the previous answer requires, while vocal and postural effort appear insensitive to the degree of misunderstanding. Importantly, the effect of semantic distance on cumulative upper-limb effort was uniformly distributed across the sample: the random-slope variance for cosine similarity was negligible across all three grouping levels (concepts, participants, and dyads), indicating that the effect did not vary systematically by performer, dyad, or concept.

### Effort does not predict communicative success

In an exploratory analysis, we asked whether the effort performers invested in repair translated into improved understanding. Across four complementary outcomes (1. whether the misunderstanding was resolved on the first attempt, 2. how semantically similar the guesser’s answer was to the target concept, 3. how quickly understanding was reached across correction attempts, and 4. whether understanding was ever achieved across the full sequence), modality and expressibility were the dominant predictors and effort was not (Fig. 1d).

Multimodal trials outperformed vocal-only trials across all outcomes. The predicted probability of reaching understanding at first attempt was 0.46 (95% CrI: [0.36, 0.55]) for multimodal and 0.18 (95% CrI: [0.06, 0.43]) for vocal-only (contrast = -0.27, 95% CrI [-0.42, -0.03]). The predicted cosine similarity of the guesser’s answer to the target concept was 0.48 (95% CrI: [0.46, 0.51]) for multimodal and 0.38 (95% CrI: [0.33, 0.43]) for vocal-only (contrast = -0.11, 95% CrI [-0.15, -0.05]). Multimodal trials also showed the lowest probability of never being resolved (P(never) = 0.25, 95% CrI: [0.19, 0.32]) compared to vocal-only trials (P(never) = 0.51, 95% CrI: [0.28, 0.73]), and the highest probability of ever being resolved (P(ever) = 0.75, 95% CrI: [0.68, 0.81]) compared to vocal-only trials (P(ever) = 0.50, 95% CrI: [0.26, 0.74]). Gesture-only performances were largely indistinguishable from multimodal in terms of success measures, although they tended in the same direction as vocal-only trials. Similarly, gesture-only performances did not reliably differ from vocal-only performances (Supplementary Material section F, Fig. 5).

Despite the clear effort increase during repair, this investment does not translate into communicative gain (see Supplementary Material section F, Fig. 10). Effort slopes were small, inconsistent in direction, and non-credible across the large majority of effort × modality combinations (see Supplementary Material section F, Table 10). A small number of exceptions ran in opposing directions and warrant cautious interpretation.

Instantaneous vocal effort predicted faster understanding in vocal-only trials and higher probability of being understood at first attempt; and greater instantaneous postural effort at first attempt predicted higher probability of eventually reaching understanding in vocal-only trials, presumably indexing speaker commitment when gesture is unavailable. In the opposite direction, higher instantaneous upper-limb effort in multimodal trials was associated with lower probability of eventual understanding, possibly marking difficulty rather than driving success.

## Discussion

When people fail to communicate a concept through non-linguistic means, they do not only reorganize their behavior – they invest more physical effort. In the present study, they did so in a specific way: not by intensifying their actions, but by doing more over time. This distinction between intensification and elaboration may point toward a principle of how the body responds to misunderstanding in a communicative context under conditions where detailed linguistic repair is unavailable. Whether the elaboration consists of a qualitatively distinct motor strategy – for instance, recruiting additional movement units or extending spatial trajectories – or whether it emerges partly from the temporal structure of repetition itself remains an open question. What our data show is that the overall effort investment increased with misunderstanding, and that this increase unfolded over time rather than in the form of amplified bursts. The increase was also flexible – sensitive to how large the misunderstanding was and shaped by the articulatory resources available. We thereby show that misunderstanding literally recruits investment of physical (and by proxy an estimate of metabolic) cost, thereby providing a fundamental grounding of “cost” that has so far remained implicit in accounts of effort and efficiency in linguistics.

These findings extend existing accounts of how environmental constraints reshape communicative behavior. Lindblom’s H&H Theory (Lindblom, 1990) predicts that a system under pressure produces a clearer, more precise signal. Rather than contradicting this prediction, our results suggest a refinement: in contexts where understanding is not hampered by environmental conditions that prevent the successful transmission of the signal itself (e.g. speech in noisy environments) but where the feedback is diagnostic – revealing not just that understanding failed but where and how (as in the form of guesses of what the communicated message meant) – performers can mount a targeted, structured repair that expands effort across time rather than amplifying individual bursts. Although repair in everyday interaction is often due to problems in hearing, which may lead to intensified repeats of the original message, the present study’s guesses equate most to targeted repair initiations, such as an interlocutor asking ‘you mean x’? (also called restricted offers in the conversation analytic literature, e.g. Dingemanse & Enfield, 2015). As such, our operationalization allows us to claim, for the first time, that performers invest more physical effort – rather than only reorganizing the content of the message – specifically when feedback helps to specify what was not understood.

Crucially, elaboration is not reflected in the same manner across dimensions. Upper-limb movement shows the greatest capacity for cumulative increase: gesture offers multiple routes through which more structure can be added such as pointing, tracing, representing, and the increase may also reflect cognitive engagement with the target, given gesture’s link to conceptual and lexical processing (Hostetter et al., 2007; Krauss & Hadar, 1999). Vocal effort showed a more modest increase, possibly reflecting the limited degrees of freedom available for non-linguistic vocalization: once referential breakdown has occurred, the semantic space is harder to navigate without the semiotic flexibility that gesture affords. Postural effort also increased cumulatively across conditions; its smaller magnitude is consistent with a stability-oriented function (Massion, 1992) and its presence even in the vocal-only condition suggests that misunderstanding imposes non-representational demands on the whole body, rooted in cognitive load or self-regulatory mechanisms (Lin et al., 2026; Yardley et al., 1999).

The diverging articulatory and meaning-making affordances in gestural and vocal modality hint that the two modalities complement each other, though our data show this complementarity is asymmetric. Vocal effort was substantially lower in the multimodal than in the vocal-only condition, indicating that the voice carries less of the communicative load when the upper limbs are available. The complementary comparison – upper-limb effort in multimodal versus gesture-only condition – pointed in the same direction (smaller upper-limb effort when voice is present) but its interval included zero. With respect to communicative success, gesture and vocalization together outperformed vocalization alone, but not gesture alone across all four outcomes. Together, this implies that adding gesture to voice helps more than adding voice to gesture, further corroborating the semiotic flexibility that gesture affords. Interestingly, we found no credible difference in communicative success between gesture-only and vocal-only trials, despite the gestural advantage reported in the literature (Fay et al., 2014; Fay et al., 2022). Unlike prior work, our model includes expressibility, which was by far the strongest driver of success (see Supplementary Material section F, Table 10). Because expressibility captures how well a concept’s affordances match a given modality, it likely absorbs the variance that previously appeared as a gesture-vocal difference – extending our earlier work on guessability in this dataset (Ćwiek et al., 2026). The modality contrast, in other words, may reflect concept-modality fit rather than an intrinsic advantage of gesture over vocalization.

A central finding of our study is that effort and success are largely unrelated in our data: although performers recruited more effort under misunderstanding, that effort did not lead to better outcomes. Such decoupling is not uncommon (Wolff et al., 2025) and the available evidence on this question is mixed. While some work hints at greater effort bringing greater success (Drijvers & Özyürek, 2017), many studies leave the relationship unresolved (Holler & Wilkin, 2011; Rasenberg et al., 2022; Trujillo et al., 2021) or fail to find it altogether (Bögels et al., 2024). Our data indicate that repair may be a context where effort serves a different function altogether. In a noisy room, a strong, clear signal is a precondition for any interaction to take place at all. Misunderstanding may call for something else. Instead of more effort, the interlocutor could benefit from change of tempo, greater informativity of individual units, or – as our data show – the number of communicative channels available. The persistent effect of expressibility on both effort and success in our data points in the same direction: a specific meaning shapes how effortful it is to communicate and how readily it is understood, somewhat independent of how hard the performer works. What does seem clear is that the communicative value of multimodality lies not in efficiency but in the richer articulatory palette it affords.

Yet effort is not decoupled from the communicative context altogether. Our second pre-registered analysis shows that the relationship between effort and understanding runs in the other direction: rather than driving the communicative outcome, effort is a response to it. Cumulative upper-limb effort scaled with the degree of misunderstanding, decreasing as the guesser’s previous answer approached the target meaning. This calibration was specific to the arms; neither vocal nor postural effort tracked the degree of misunderstanding. It was also uniform across the sample, varying little across performers, dyads, or concepts, which suggests that the adjustment of arm effort to common ground operates similarly across individuals rather than reflecting idiosyncratic strategy. Similarly to the correction effect in H1, instantaneous measures were largely unmoved by misunderstanding. The consistent picture is that misunderstanding reshapes how long and how much performers communicate, not how forcefully. This is in support of audience design accounts (Clark & Murphy, 1982; Horton & Keysar, 1996) proposing that people monitor the current state of shared understanding and tune their behavior to their partners when necessary. Future work shall determine the qualitative nature of this attunement, exploring the relationship between different types of misunderstanding and the changes they trigger.

Our findings must be interpreted in light of several limitations. The controlled design of our paradigm, while necessary for isolating effort modulation, comes with trade-offs in ecological validity. Whole-body movement without speech amplifies physical demands beyond what typical face-to-face interaction requires, and in naturalistic settings linguistic structure carries much of the repair work. We view this as a necessary first step: charade-like referential tasks share fundamental properties with linguistic interaction: they are inherently communicative, meaningful, and intersubjective (Christiansen & Chater, 2022). Identifying effort modulation in this simplified context establishes a foundation for investigating these phenomena in more naturalistic settings.

A related limitation concerns what counts as communicative success. Our design treats understanding as an exact match between the guesser’s answer and the target concept, whereas real interaction typically requires only sufficient mutual understanding to support the next action. More consequentially, our paradigm places most of the repair work on the performer: the guesser contributes only a single guess per performance, leaving little room to act on the performer’s increased effort. Research on referential tasks with freer interaction suggests that more distributed repair (where partners jointly signal and respond to misunderstanding) may be what drives communicative success (Rasenberg et al., 2022). It is therefore possible that effort does not help in our paradigm precisely because the partner who could benefit from it has limited room to act on it.

A second set of limitations concerns the measurement of effort itself. The three dimensions we selected – arm torque change, acoustic amplitude envelope, and change in centre of pressure – capture general biomechanical demands on the body, but they are necessarily selective: effort may also be expressed through articulators and movement qualities our measures do not index. The substantial concept-level variance in our models hints that concepts may also differ in which acoustic-motor components they draw on to convey meaning, and thus in the kind of effort they recruit. While our population-level effects suggest that the selected features capture a general mechanism, future work should examine other articulators (e.g. head or facial expressions) and whether other dimensions of movement such as speed, spatial precision, or inter-articulator coordination carry additional information that our measures do not. In particular, the coordinative cost of orchestrating multiple articulators simultaneously remains unaddressed here and may represent a distinct dimension of multimodal effort with its own scaling properties. Our preregistration (Kadavá et al., 2025) includes an example of such exploratory analysis, which we will address in an upcoming work. Taken together, these limitations point toward productive directions for future research without undermining the core finding: that the body responds to misunderstanding with cumulative, distributed effort reliably, across modalities, and, for the arms, in proportion to the degree of communicative failure.

Together, the absence of an effort-success link and the responsiveness of effort to feedback point toward a reframing of what communicative effort represents. Rather than a resource allocated toward success, effort appears to function as a continuous index of the interaction’s epistemic state: signaling bodily mobilization when understanding breaks down and scaling with the distance from common ground. In this respect it resembles sensorimotor communication, where performers modulate their actions to move away for their baseline which then stands out as a signal (Pezzulo et al., 2019). These findings sit in tension with accounts of principle of least effort and effort aversion (André et al., 2026; Job et al., 2024). If people generally avoid effortful behavior (David et al., 2024), why would they increase physical investment in response to misunderstanding without clear returns in success? The standard evolutionary view holds that efficient signals convey as much as possible at minimal cost but our findings indicate a complementary pressure: effort may be organized not around maximizing informational yield but around sustaining the communicative cycle itself. What costs something is not saying more but staying in the exchange long enough to be understood. Research on challenging tasks suggests that this kind of investment is not irrational: effort carries intrinsic social value, enhancing the perceived worth of its product including fostering social belonging (Inzlicht et al., 2018). On this view, the effort enhancement we observe may be less about solving the problem and more about affirming commitment to the partner at the moment the interaction is most at risk of failing.

The relational function of effort may extend beyond humans. When animals repeat and vary their signals in the face of failure, this is taken as one of the objective criteria of intentional communication (Townsend et al., 2017). Our findings suggest that intentionality may leave a measurable physical trace, as sustained effort as operationalized here must carry a metabolic cost. What happens to that trace when breakdown escalates into conflict, when the interaction is particularly vulnerable, or when competing demands leave fewer bodily resources available? These are the conditions where the effort to stay in it together – for humans and animals alike – matters most.

## Methods

### Preregistration and ethics

The experimental design was preregistered at https://osf.io/3nygq on December 13, 2023 prior to data collection; and the hypotheses and analyses were preregistered at https://osf.io/8ajsg/overview on April, 11 2025 prior to data analysis, using pilot dyad. We report hypotheses, predictions, outcome variables and analyses as preregistered unless otherwise indicated. All preregistered scripts are available at a frozen repository (https://github.com/sarkadava/FLESH_Effort/tree/preregistered) and changes between the pre-registered and final processing pipelines are documented in a dedicated change-tracking report, which provides a line-by-line comparison of the two versions. Any deviations reflect improvements to data quality and processing robustness rather than changes motivated by the results, and are individually motivated in the Supplementary Material. All participants provided informed consent and were reimbursed by course credit or monetary reward. This study was approved by the Ethics Committee Social Sciences (ECSS) of Radboud University (reference number 22N.004687). All materials, data and code are publicly available on Github, with reproducible analysis code provided in the electronic Supplementary Material (https://sarkadava.github.io/FLESH_Effort/).

### Participants

Participants were 144 native Dutch speakers (mostly bilingual with a high proficiency in English) recruited from a Dutch university for course credit or monetary reward. They were able to sign up as a team or alone through the university recruitment system. We collected data about their personality traits (Denissen et al., 2008), handedness (Oldfield, 1971), and demographics. Additionally, we asked to assess how familiar and comfortable they feel with their experimental partner. In total, we collected data from 72 dyads, a number covering for exclusions (outlined below) and based on pre-registered power analysis.

After exclusion, our pool of participants consisted of 61 dyads (i.e. 122 participants). Of these, 99 were females, 20 were males, 1 non-binary, 1 other and 1 preferred not to say, with a mean age of 19.9 years (SD = 1.9, range = 17–27). The majority were right-handed (108), with 14 left-handed participants. The final count for trials was 1244 for the gesture condition, 1933 for the vocal condition, and 1251 for the multimodal condition.

### Data exclusions

Before data processing, we excluded one dyad due to consent withdrawal. During data processing, we excluded nine dyads due to recording errors. Further, following preregistered exclusion criteria, we removed one dyad due to persistent high residual error (larger than 25 mm) in pose estimation. Additionally, we excluded 628 trials that had high error for inverse kinematics (larger than 40 mm). Lastly, we excluded 165 trials in which participants violated the condition rule (e.g. not using both modalities in the combined condition, using sound in the gesture condition, etc.) to ensure that effort is strictly unimodal or multimodal, and 12 trials in which participants used speech for concept explanation.

## Materials

### Lab equipment

#### Cameras dedicated for motion tracking with computer vision

For video recording, we used three Elgato Facecam cameras located at a movable arch. The Elgato settings were set at a relatively low resolution of 940x540 to increase mobility of the data, a shutter speed of 1/200s to decrease motion blur increasing motion tracking performance (and an ISO 354, to stabilize contrast due to lower exposure time). We used a custom Python script to capture and write videos, using the opencv and ffmpegcv packages (Kadavá et al., 2024). We write the videos in raw video codec with a frame rate of 60 fps. After each session, we compressed each video to the XVID codec. At the beginning of each session, we recorded 1-minute calibration videos with a checkerboard and charuco board, which allowed us to estimate the intrinsic and extrinsic angles of the cameras needed for 3D pose estimation (Karashchuk et al., 2021; Matthis et al., 2022; Pagnon et al., 2022a). The experimenter followed a protocol for uniform camera calibration (e.g. covering a vertical plane with small circular motions with the outermost corners defined by the edges of the balance board).

#### Balance board

For analysis of postural adjustments, we used a balance board that has been designed by the Technical Support Group of Donders Institute. The board incorporated sensors adapted from the Wii Balance Board, which have been redesigned. The system was synchronized in time with an accuracy of one millisecond and in space with an accuracy of several submillimeters. An A/D conversion was performed by a National Instruments card, specifically the USB-62221, which was connected to the PC via USB. This card collected four signals at a sampling rate of 400 Hz.

#### Audio recording

For audio recording, we used a C520 head-mounted condenser microphone, with a low-noise 2-channel DAP PRE-202 microphone amplifier (with +48 V phantom power and low-cut function turned on) of which the gain was set at a constant level (25%) over the entire experiment. The volume level in the computer’s sound panel setting was set to 100. The audio signal was split using an XLR splitter, allowing us to record the audio signal at 16 kHz as an LSL stream transmitted via a Linux-based Minux system and a higher quality 48 kHz via another recording PC using Audacity. The 48 kHz audio recording was used for extraction of acoustic features (e.g. fundamental frequency, amplitude envelope, center of gravity, etc.) Additionally, the audio signal leading to this PC was split once more, allowing us to connect noise-canceling Sony headphones (WH-1000XM5). These were used by the guesser. The motivation was to make sure that the audio signal from the performer could be smoothly perceived by the guesser, since a black curtain separating the two participants hindered the intensity of the audio signal in the room. The guesser also used a second pair of identical headphones in gesture-only condition: these did not transmit any audio signal as they were intended to ensure that the performances were perceived only via visual modality.

#### One-way screen

We used a one-way screen that has been designed by the Technical Support Group at the Max Planck Institute for Psycholinguistics. The motivation for the one-sided view was to limit any spill-over feedback or communication from the guesser through gestures or facial expressions. This enabled maximal control of the communicative flow and feedback while sustaining the co-presence of two participants in one room.

#### Experiment software

The experiment was managed by a custom Python script, using the PsychoPy package, and a buttonbox that was implemented in the script with the RuSocSci package. It outputed a CSV file that stored information about accuracy for each trial (i.e. concept).

All specifications regarding laboratory equipment are described in detail in the method preregistration. All scripts for recording are available at https://github.com/sarkadava/FLESH_Effort/tree/main/00_labscripts.

### Stimuli

The stimuli were selected from a list of 206 concepts. This list included 100 Leipzig-Jakarta List concepts (Tadmor et al., 2010) reflecting concepts that are common across a large range of languages and are most resistant to borrowing from other languages. Additionally, 100 concepts were included varying in sensory expressibility (Lynott et al., 2020; Lynott & Connell, 2009). In a previous experiment (Ćwiek et al., 2026), 227 native Dutch participants rated the current concepts on a continuous scale for how well they could be communicated without language across three modalities: gesture, vocal, and combined. From the 206-item list, we excluded concepts with any expressibility value below a threshold (mean expressibility – 1 SD) to avoid low-expressible concepts. Using a custom Python script, we constructed three modality-specific 28-item lists of top-ranked expressible concepts, ensuring no overlaps between them. Body-related concepts (e.g. ‘tooth,’ ‘ear,’ ‘tongue’) were excluded to prevent indexical reference to the body itself. They were replaced with taste-related concepts that are generally low in expressibility, aligning with secondary research interests. The final 84-item list maintains expressibility statistics comparable to the original list. The final concepts included nouns, verbs and adjectives. Stimuli lists were pre-randomized to ensure balanced occurrences across sessions (each concept appears at least 10 times, or 12 if additional dyads are included). However, the occurrences are not precisely balanced due to a processing mistake.

### Procedure

Participants engaged in a referential, charade-like game in a lab-based setting in which one person (the performer) expresses a concept without using language, and the other (the guesser) guesses its meaning. This paradigm has been widely used to explore the process of grounding the meaning of novel expressions built on various modalities such as drawing (Hawkins et al., 2023), gestures and vocalizations (Ćwiek et al., 2021; Fay et al., 2014; Fay et al., 2022; Macuch Silva et al., 2020), or other, more artificial systems (Selten & Warglien, 2007). Importantly, we adapted the referential game to include feedback. After each guesser’s answer, the performer saw the exact response, and if it did not match the target concept, they had to repair their original performance. Each participant performed novel expressions of 21 concepts, for a total of 42 concepts per dyad, across three within-subject modality conditions: vocal, gesture, and combined (i.e. 7 concepts per condition). This design allowed us to assess how effort is affected by and distributed across modalities. The order of modality conditions was randomized. Each modality was introduced with instructions, followed by two practice concepts and seven trials. After each performance, the guesser typed their answer on a keyboard. If correct, they proceeded to the next concept. If incorrect, both saw the incorrect response, and the performer repeated the concept while the guesser made another attempt. Up to two repair attempts were allowed before moving on. For our purposes, the first performance of a concept established a baseline, and the potential repairs that followed are what we called first and second corrections. Correctness was judged by the experimenter, who checked only for typos; synonyms were not accepted to ensure consistency across sessions. Participants switched roles within and between conditions.

Instructions, stimuli, and feedback were presented on a screen via a custom PsychoPy script. Performers indicated the start and end of each production with a predefined gesture, crossing their arms in front of their body. The guesser was seated at a table with a computer screen to present instructions and feedback. They used a keyboard to type in the answers, which was possible only after the performers indicated the end of a production.

All specifications regarding experimental setup are described in detail in the method preregistration.

## Data processing

### Pre-processing

Each session produced several recorded streams (i.e. video frame stream, balance board stream, and audio stream), which was read and processed via custom Python scripts. These streams were natively synchronized using LabStreamingLayer (Kothe et al., 2025), which captured all streams in an XDF formatted file. We used buttonbox timestamps to isolate trial segments in each stream and associate them with metadata (e.g. condition, participant number, etc.). For each trial, the output of interest includes a 60 fps video, 48 kHz audio, and balance board data. The audio and video were visually inspected to ensure they start and end when the performer signals the beginning and the end. Trials that started late or ended earlier were adjusted.

### Motion tracking

We used OpenPose (Cao et al., 2019), specifically its 135-marker model, to obtain a 2D skeleton with 135 body keypoints (per camera), including hands and face, with a sampling rate of 60 Hz.

To convert multiple 2D data streams into 3D positional data, we used triangulation based on calibrated cameras. Triangulation was performed using Pose2Sim (version 0.10.39, Pagnon et al., 2022b). To triangulate all 2D skeleton data, we first calibrated the cameras using a checkerboard to determine the intrinsic and extrinsic parameters (Supplementary Material section B, Fig.1–2). For intrinsic calibration, we obtained an error of 0.24 pixels for each camera (recommended below 0.5 pixels). Residual (extrinsic) calibration errors across all cameras were 0.96 cm. The mean reprojection error for all points across all frames and trials was 1.49 cm (values below 1 cm are recommended, but up to 2.5 cm is acceptable). The triangulated data were directly smoothed within the Pose2sim pipeline, which was set as a 4th-order, 10Hz low-pass, zero-phase Butterworth filter; however, this still yielded jittery trajectories, so we smoothed the coordinates further with a Savitzky-Golay 3rd-order polynomial filter with a span of 167 ms which is useful for removing high-frequency jitter. Before obtaining inverse kinematics, we further processed the data to align coordinates with OpenSim requirements (e.g. the feet need to touch the floor).

We further used Pose2sim’s implementation to scale a skeletal model to each participant’s body (i.e. height and weight). The values for weighting the importance of each keypoint were kept at default values. We then used the scaled model to calculate joint angles, i.e. angles between the line of the proximal and distal segment of a joint, for each trial (represented by the coordinates obtained in the previous step).

Joint angles were then used to obtain generalized net joint torques via the OpenSim package (Seth et al., 2018). Joint torque is a measure of rotatory force with which a segment moves, and can be estimated by inverse dynamic analysis. To prevent amplification of noise when solving inverse dynamics, we first smoothed the joint-angle data using a Savitzky-Golay filter with a span of 560 ms and a polynomial order of 3. The average root mean square error was 0.03 m.

### Processing of key time series

#### Amplitude envelope

We used the high-sampling 48 kHz audio data to extract the amplitude envelope of the acoustic signal. We followed a method by Tilsen and Arvaniti (Tilsen & Arvaniti, 2013), implemented in Python by Pouw (Pouw, 2024). We used a bandpass and a 2nd-order, 10Hz low-pass, zero-phase Butterworth filter. Finally, the data were normalized to the range 0 to 1 within each participant.

#### Joint torque change

Joint torque data were smoothed with a Savitzky-Golay filter with a span of 560 ms and polynomial order of 1. Further, we obtained the time derivative of the joint forces, namely, the torque change (in Nm/s). We smoothed the torque change with a Savitzky-Golay filter with a span of 560 ms and polynomial order of 1. To create an aggregated derivative measure for the whole arm, we computed an Euclidean sum over the torque change of all key points belonging to the arm.

#### Change in center of pressure

We computed the change in 2D magnitude in the center of pressure and smoothed it using the Savitzky-Golay filter with a span of 102 ms and polynomial order of 5.

All time series were merged on a common sampling rate of 500 Hz.

### Extraction of key variables

From the processed time series, we derived six effort variables across three channels listed below. To fully capture the effort dynamics, we considered both cumulative and instantaneous effort for all these three measures. Cumulative effort reflects the total exertion over time, which is particularly relevant for sustained actions and overall physical demand. We quantified this by integrating each feature across time, providing a measure of accumulated intensity, torque change, and postural shifts. Instantaneous effort, on the other hand, highlights peak exertion at specific moments. By analyzing peak values of the signals, we can identify the local physical demands on the body. This dual approach allowed us to differentiate between continuous exertion and sudden bursts of effort, offering a more comprehensive perspective on physical cost across communicative attempts.

#### Amplitude envelope (vocal effort)

For vocal effort, we computed the cumulative and instantaneous features of the amplitude envelope as a continuous index of respiratory(-vocal) engagement. We opted for intensity over f0 or formants because non-linguistic vocalizations do not always contain phonation, making spectral measures unreliable across our stimulus set.

#### Torque change (upper-limb effort)

For upper-limb effort, we computed cumulative and instantaneous features of aggregated net joint torque change, which accounts for segment masses the inertial properties of moving limb segments.

#### Change in center of pressure (postural effort)

For postural effort, we computed cumulative and instantaneous features of change in the center of pressure (COPc), capturing fluctuations in postural control, which will be very much driven by upper-limb movement and whole-body engagement when present.

#### Degree of misunderstanding

We operationalized a degree of misunderstanding as a semantic distance between the guesser’s answer and the performer’s target concept, computed as the cosine similarity of the ConceptNet word embeddings (Speer et al., 2018). For each target–answer pair from the experiment, we computed cosine similarity using Dutch word embeddings from ConceptNet (numberbatch version 19.08, see preregistration for validation). For pairs not represented in the embeddings (e.g. two-word answers), we conducted an online rating study, collecting data from 16 Dutch native speakers who were asked to rate how similar the word pairs felt.

## Data analysis

### Confirmatory analysis

Confirmatory analyses concern pre-registered analysis and hypotheses. All statistics were performed using R (Team, 2025). We fitted Bayesian mixed effects models, using the brms package (Bürkner, 2017), to test two preregistered hypotheses:

*H1: Correction recruits more physical effort than the baseline performance*.
*H2: A higher degree of misunderstanding will require a performer to engage in more effortful correction*.

For H1, we fitted six sets of models for the six investigated dependent variables: 1) arm torque integral (cumulative upper-limb effort), 2) envelope integral (cumulative vocal effort), 3) COPc integral (cumulative postural effort), 4) arm torque peak mean (instantenous upper-limb effort), 5) envelope peak mean (instantenous vocal effort), 6) COPc peak mean (instantenous postural effort). The primary predictor of interest was correction. To identify the most appropriate model specification, we fitted four variants for each dependent variable, increasing in complexity.

Model 0 included only the primary predictor as a fixed effect.

Model 1 additionally included covariates: familiarity between the guesser and the performer, extraversion of the performer (BFI), expressibility of the concept, modality, and trial number. These were included not as predictors of primary interest but as theoretically motivated adjustment variables. To guide covariate selection, we constructed a directed acyclic graph (DAG) reported in full in the Supplementary Material (section E). The DAG formalized our assumptions about the causal structure of the data – specifically, which variables influence both the likelihood of receiving a correction and the effort expended during it. A variable that affects both the treatment (correction) and the outcome (effort) without lying on the causal path between them constitutes a confounder; failing to adjust for it would bias estimates of the correction effect. In our DAG, familiarity, extraversion, expressibility, modality, and trial number all plausibly met this criterion: for example, more expressive concepts are easier to communicate and therefore less likely to require correction, while also requiring less physical effort. Leaving expressibility uncontrolled would therefore inflate the apparent effect of correction on effort. All fixed effect predictors were pre-registered before data analysis. Model 1 included random intercepts and slopes for all predictors that vary within each grouping factor (participants, dyads, concepts). Although dyad-level random effects were not formally pre-registered, we included them because participants were nested within dyads by design – dyad partners shared the same interactional context, and ignoring this clustering would lead to underestimation of uncertainty in fixed-effects estimates.

Model 2 extended Model 1 by estimating the full covariance structure between random intercepts and slopes for each grouping factor.

Model 3 was identical to Model 2 but did not estimate correlations between random effects. This simplification was motivated empirically: in Model 2, most correlation parameters had credible intervals spanning the full [-1, 1] range, indicating the data did not contain sufficient information to estimate them reliably, and their inclusion caused sampling inefficiency (low effective sample sizes).

The reporting model was selected from Models 1–3 based on convergence quality (Rhat < 1.01, bulk ESS > 1000) and, among converged models, predictive performance (ELPD-LOO). All models were fit with weakly informative priors that were unbiased with respect to H0 and H1. All categorical covariates were contrast-coded, and all continuous covariates were z-scored or centered.

Model 4 extended the best converging model from Models 1–3 by adding interaction terms between the primary predictor and theoretically motivated covariates. Specifically, the fixed effects included: correction × modality (testing whether effort escalation at corrections differs by communicative channel), correction × expressibility (testing whether the correction effect depends on concept expressibility), correction × familiarity (testing whether familiar dyads enhance effort differently at corrections), correction × extraversion (testing whether personality modulates communicative effort responsiveness), and modality × expressibility (testing whether the expressibility-effort relationship is modality-specific). These interactions were pre-registered. The random-effects structure retained the specification of the best converging base model. Model 4 was fitted for exploratory purposes to identify theoretically motivated modulations of the correction effect, and results are interpreted accordingly.

For H2, we selected the best-performing model from H1 for each dependent variable and extended it with two additional predictors: the cosine similarity of the previous answer to the target concept, and the effort expended in the preceding communicative attempt. Previous effort was included as a statistical control: effort levels may vary across trials due to differences between concepts, dyads, and participants. Including previous effort accounts for this trial-level continuity, ensuring that any effect of answer similarity on current effort reflects genuine calibration to the previous answer rather than between-trial differences in effort. Note that this is a deviation from the preregistered analysis, where we intended to model change in effort directly. However, effort change scores – computed as the difference between consecutive attempts – produced distributions with substantial positive and negative values that proved difficult to model reliably, explaining less than 5% across all six dependent variables. Modeling current effort with previous effort as a covariate recovered the same inferential target while avoiding these estimation problems. The preregistered change-score models are reported in full in the Supplementary Material (section E) for transparency.

Across the four models we fitted (Models 1–4), we adopted the model comparison framework articulated in McElreath (McElreath, 2018): rather than treating model fitting as a selection problem in which one model is chosen and the rest discarded, we treated the set of models as complementary lenses on the same data, each estimating a different combination of parameters. Model 3 served as our primary reporting model, on the grounds described above (best balance of convergence diagnostics and ELPD-LOO among Models 1–3). Estimates of the primary effects were stable across Models 1–4, indicating that our inferences did not depend on a particular random-effects specification. Where parameters of interest were not estimated in Model 3, we drew on the models that did: Model 2 for correlations between random intercepts and slopes, and Model 4 for theoretically motivated interactions.

### Exploratory analysis

Beyond the preregistered analyses, we also investigated questions that emerged after fitting the models of primary interest. In the first set of exploratory models, we asked whether cumulative effort can be explained simply by duration – that is, whether more time merely accumulates more movement – or whether it also reflects modulation of amplitude. Trial duration was fitted as one additional predictor for each of the best-performing models from H1 for each cumulative dependent variable.

In the second set of exploratory models, we asked whether the effort dedicated to a correction depends on the participant’s average baseline level of effort at first performance. We fitted the best-performing models from H1 for each dependent variable with an additional predictor: the effort of the first performance, entered in interaction with the correction predictor. This allowed us to test whether correction-related effort escalation is uniform across performers or modulated by their initial effort level.

In the third set of exploratory models, we asked whether effort invested across the correction sequence predicts communicative success. We examined four complementary outcomes: whether misunderstanding was resolved on the first attempt (Bernoulli model), how semantically similar the guesser’s following answer was to the target concept (zero-one inflated Beta model), how quickly understanding was reached across the full sequence (cumulative ordinal model), and whether understanding was ever achieved (Bernoulli model). For each outcome, effort predictors – arm torque change, amplitude envelope, and COPc, both cumulative and instantaneous – were entered alongside modality as interaction terms, allowing us to test whether the effort-success relationship differs by communicative channel. All effort predictors were log-transformed and centered on the active-channel mean; in inactive channels, values were attenuated by a constant factor to reflect noise rather than signal.

## Acknowledgments

We would like to thank all participants of this study. Special thanks also go to the Donders lab coordinator Jiska Koemans and the Donders research integrity officer Miriam Kos. We are especially grateful to the members of Donders Technical Support Group, namely Erik van den Berge, Norbert Hermesdorf, Gerard van Oijen, Maarten Snellen and Pascal de Water, for their invaluable help with the technical setup. Finally, we thank the student assistants and interns in project FLESH – Jet Lambers, Justin Snelders, Gillian Rosenberg, Hamza Nalbantoğlu – who supported this project through their efforts in participant recruitment, data collection, data processing, and annotation.

## CRediT author statement

ŠK: conceptualization, methodology, software, formal analysis, investigation, writing – original draft, writing – review & editing, visualization; AĆ, SF and WP: conceptualization, methodology, supervision, writing – review & editing, project administration, funding acquisition; JH: methodology, supervision, writing – review & editing. All authors approved the final paper.

## AI statement

Generative AI models were used for two areas: code review/debugging (Mistral) and text revision (Claude). The use of generative models was in accordance with the guidelines proposed by the Deutsche Forschungsgemeinschaft (German Research Foundation): https://www.dfg.de/en/service/press/press-releases/2023/press-release-no-39

